# BioLaboro: A bioinformatics system for detecting molecular assay signature erosion and designing new assays in response to emerging and reemerging pathogens

**DOI:** 10.1101/2020.04.08.031963

**Authors:** Mitchell Holland, Daniel Negrón, Shane Mitchell, Nate Dellinger, Mychal Ivancich, Tyler Barrus, Sterling Thomas, Katharine W. Jennings, Bruce Goodwin, Shanmuga Sozhamannan

## Abstract

**Background:** Emerging and reemerging infectious diseases such as the novel Coronavirus disease, COVID-19 and Ebola pose a significant threat to global society and test the public health community’s preparedness to rapidly respond to an outbreak with effective diagnostics and therapeutics. Recent advances in next generation sequencing technologies enable rapid generation of pathogen genome sequence data, within 24 hours of obtaining a sample in some instances. With these data, one can quickly evaluate the effectiveness of existing diagnostics and therapeutics using *in silico* approaches. The propensity of some viruses to rapidly accumulate mutations can lead to the failure of molecular detection assays creating the need for redesigned or newly designed assays.

**Results:** Here we describe a bioinformatics system named BioLaboro to identify signature regions in a given pathogen genome, design PCR assays targeting those regions, and then test the PCR assays *in silico* to determine their sensitivity and specificity. We demonstrate BioLaboro with two use cases: Bombali Ebolavirus (BOMV) and the novel Coronavirus 2019 (SARS-CoV-2). For the BOMV, we analyzed 30 currently available real-time reverse transcription-PCR assays against the three available complete genome sequences of BOMV. Only two met our *in silico* criteria for successful detection and neither had perfect matches to the primer/probe sequences. We designed five new primer sets against BOMV signatures and all had true positive hits to the three BOMV genomes and no false positive hits to any other sequence. Four assays are closely clustered in the nucleoprotein gene and one is located in the glycoprotein gene. Similarly, for the SARS-CoV-2, we designed five highly specific primer sets that hit all 145 whole genomes (available as of February 28, 2020) and none of the near neighbors.

**Conclusions:** Here we applied BioLaboro in two real-world use cases to demonstrate its capability; 1) to identify signature regions, 2) to assess the efficacy of existing PCR assays to detect pathogens as they evolve over time, and 3) to design new assays with perfect *in silico* detection accuracy, all within hours, for further development and deployment. BioLaboro is designed with a user-friendly graphical user interface for biologists with limited bioinformatics experience.

## Background

Emerging and reemerging infectious diseases have serious adverse impacts on society with respect to lives lost and annual economic losses [1-5], as being witnessed currently in the COVID-19 outbreak caused by a novel Coronavirus, SARS-CoV-2. As of March 9, 2020, there have been 113,584 confirmed cases of COVID-19 around the globe and 3,996 deaths [6-8]. Factors such as climate change, urbanization, zoonotic spillover, international travel, lack or breakdown of public health care systems, natural or man-made disasters, and pathogen evolution to name a few, contribute to infectious disease emergence and sustainment [2].

A unique challenge associated with emerging infectious diseases is rapidly identifying the etiological agent and developing or repurposing existing medical countermeasures (MCMs), including diagnostic assays, to curtail the spread of an outbreak. With respect to reemerging infectious diseases the challenge is to determine whether existing MCMs are effective or not, and in the latter case, develop or modify the MCMs in a timely manner. In some cases, the failure of existing MCMs can be accounted for by genetic drift and shift and the resulting altered genotypic and phenotypic profiles of the newly emergent pathogen. Ebolavirus is one such reemerging infectious disease agent that exhibits high degrees of genetic changes in every new outbreak.

Ebola virus disease (EVD) is one of the deadliest infectious diseases that continues to plague Central and Western Africa. It was discovered in 1976 when two consecutive outbreaks of fatal hemorrhagic fever occurred in different parts of Central Africa [9]. The first outbreak occurred in the Democratic Republic of the Congo (DRC, formerly Zaire) in a village near the Ebola River, which gave the virus its name [9]. Since 1976 more than 25 outbreaks have been recorded. The average case fatality ratio (CFR) for EVD is around 50% and varies from 25% to 90% in past outbreaks [9]. From 2014 to 2016, the world witnessed the worst-yet EVD outbreak, which originated in Western Africa. That outbreak started in a rural setting of southeastern Guinea, and quickly spread like a wildfire to urban areas and across borders to the neighboring countries of Liberia and Sierra Leone [10]. In the end, the outbreak infected 28,616 people and killed 11,310 of the victims (CFR of approximately 39.5%) [11]. In the current Democratic Republic of the Congo outbreak that began in August 2018, as of March 4, 2020, a total of 3,444 cases, total deaths 2,264 and 1,169 survivors have been reported [12].

Until recently, there was no approved vaccine or treatment for EVD although four experimental therapeutics, (Regeneron’s monoclonal antibody REGN-EB3, mAb-114 [Ridgeback Biotherapeutics], Remdesivir GS-5734 [Gilead] and ZMapp [MappBio Pharmaceutical] underwent clinical trials (the Pamoja Tulinde Maisha [PALM] study) during the current DRC outbreak [13]. On October 17, 2019, the Committee for Medicinal Products for Human Use granted a “conditional marketing authorization” for the Ebola Zaire vaccine ERVEBO by Merck Sharp and Dohme [14]. ERVEBO was reviewed under the European Medicines Agency’s (EMA) accelerated assessment program. ERVEBO (v290) (Ebola Zaire Vaccine (rVSVΔG-ZEBOV-GP, live) is a genetically engineered, replication competent, attenuated live vaccine that received approval from the U. S. Food and Drug Administration (FDA) on December 19, 2019 [15].

Ebolaviruses are one of three genera of filoviruses belonging to the Family *Filoviridae,* Cuevavirus and Marburgvirus being the other two. Filoviruses are non-segmented, negative-sense, single-stranded RNA viruses. Six species of Ebolaviruses have been described to date, Zaire (EBOV), Bundibugyo (BDBV), Sudan (SUDV), Taï Forest (TAFV), Reston (RESTV) and Bombali (BOMV). The prototypic viruses - EBOV, BDBV, SUDV and TAFV - have been associated with disease in humans [16, 17]. The RESTV causes disease in nonhuman primates and pigs [18, 19]. In a 2018 wildlife survey, BOMV RNA was recovered from insectivorous bats, but there are no known cases of human or animal disease caused by this species [20].

Research on EVD focuses on finding the virus’ natural reservoirs and hosts from which spill over occurs, developing preventive measures such as vaccines to protect at-risk populations, and discovering therapies to improve treatment of infections. Biosurveillance data suggested that the reservoirs of ebolaviruses may be fruit and insect eating bats [21-24]. The new BOMV was identified in a biosurveillance project conducted in Sierra Leone to identify hosts of EBOV as well as any additional filoviruses that might be circulating in wildlife [20]. Oral and rectal swabs were collected from 535 animals (244 bats, 46 rodents, 240 dogs, 5 cats) from 20 locations in Sierra Leone in 2016. Of the 1,278 samples, five samples from three Little free-tailed bats and one Angolan free-tailed bat contained ebolavirus sequences. Two full genome sequences were assembled from two of the samples and they showed nucleotide identity of 55–59% and amino acid identity of 64–72% to other ebolaviruses [20]. Based on phylogenetic analyses of sequence data, it was determined that the genome was sufficiently distinct to represent the prototypic strain of a new species, Bombali Ebolavirus (BOMV) [20].

I*n vitro* studies of the BOMV demonstrated that a recombinant vesicular stomatitis virus (rVSV) encoding the BOMV glycoprotein (GP) gene mediated virus entry into human host cells |20|. Entry and infection of rVSV–BOMV GP was also completely dependent on Niemann-Pick C1 (NPC1) protein, providing additional evidence that this is a universal receptor for filoviruses. Although not conclusive, these data indicated the potential for BOMV to infect humans. Given the high divergence from other Ebolaviruses, we examined whether existing real-time reverse-transcription PCR (rRT-PCR) assays for Ebolaviruses would detect the new BOMV.

Potential determinants of Ebolavirus pathogenicity in humans were identified by analyzing the differentially conserved amino acid positions called specificity determining positions (SDPs) between human pathogenic ebolaviruses and the non-pathogenic Reston virus [25]. Recently, this study was extended to include BOMV to assess its pathogenicity to humans [26]. At SDPs, BOMV shared the majority of amino acids (63.25%) with the human pathogenic Ebolaviruses. However, for two SDPs in viral protein 24 (VP24), which may be critical for the lack of Reston virus human pathogenicity, the BOMV amino acids match those of Reston virus. Thus, BOMV may not be pathogenic in humans [25, 26]. Nonetheless, rRT-PCR assays are important for biosurveillance of BOMV and other potential new variants in the wild.

While our study on the BOMV use case for assay design was ongoing, the world witnessed the emergence of a novel respiratory pneumonia disease (COVID-19) epidemic from Wuhan city, Hubei province, China. In the aftermath of its rapid global spread and devastating impact, on March 11, 2020, the WHO declared that COVID-19 can be characterized a pandemic threat [27]. COVID-19 was determined to be caused by a severe acute respiratory syndrome (SARS)-like corona virus (SARS-CoV-2) that quickly spread within Wuhan [28, 29] and crossed borders to >104 countries/territories/areas as of March 09, 2020 [7, 8, 30].

Using next generation sequencing, the whole genome sequence (WGS) of SARS-CoV-2 are continuously being released and shared (306 complete genomes as of March 09, 2020) with the entire research community through Global Initiative on Sharing All Influenza Data (GISAID) [31]. The release of WGS allowed us to test the BioLaboro pipeline (described in this study) to evaluate currently used diagnostic assays and to rapidly design new assays.

In a previous study, we described a bioinformatics tool called PSET (PCR signature erosion tool) and used it to show *in silico*, confirmed with wet lab work, the effectiveness of existing Ebolavirus diagnostic assays against a large number of sequences available at that time [32]. The phrase “signature erosion” used here signifies potential false-positive or false-negative results in PCR assays due to mutations in the primers, probe, or amplicon target sequences (PCR signatures). Signature erosion could also mean failure of medical countermeasures; for example, a change in the genomic sequence resulting in an amino acid change that could potentially alter the efficacy of sequence-based therapeutics [33, 34].

In this study, we describe an expanded bioinformatics pipeline called BioLaboro in which we have integrated several tools: BioVelocity®, Primer3 and PSET for end-to-end analysis of outbreak pathogen genome sequences to evaluate existing PCR assay efficacy against the new sequences, and to identify unique signature regions (BioVelocity), design PCR assays to these regions (Primer3), and test the new assays’ efficacy (PSET) against current National Center for Biotechnology Information Basic Local Alignment Search Tool (BLAST) standard databases [35]. We have used the BOMV and SARS-CoV-2 sequences as proof of concept for BioLaboro to develop and evaluate new PCR assays *in silico.*

## Results

### BioLaboro architecture

BioLaboro is comprised of three algorithms - BioVelocity, Primer3, and PSET - which are built into a pipeline for *user-friendly* applications. The user has the option to launch one of four different job types: Signature Discovery, Score Assay Targets, Validate Assay, or New Assay Discovery. Each of the three algorithms can be run individually or together as a complete end-to-end pipeline (Figure 1). For the BOMV use case, in the first phase of the pipeline BioVelocity was used to analyze a set of genome sequences for unique regions that are both conserved and signature to the target sequences selected. This was achieved by splitting a chosen representative whole genome sequence into sliding 50 base pairs (bps) k-mers. Each k-mer was then scanned against all target sequences to determine conservation. Conserved k-mers were then elongated based on overlaps and formed into contigs. These contigs were then split into k-mers ≤ 250 bps and scanned against all non-target sequences to determine specificity. All passing sequences were then elongated based on overlaps and the signature contigs were passed to the next step in the pipeline. Primer3 was then used to evaluate the signature contigs to identify suitable primers and probes for assay development. Primer3 was run in parallel against all signatures and the output was ranked by penalty score in ascending order. The top five best primer sets were passed along to the final step in the pipeline, PSET. In this step the primer sets were run through a bioinformatics pipeline which aligned the sequences against large public sequence databases from NCBI using BLAST and GLSEARCH [36] to determine how well each assay correctly aligned to all target sequences while excluding off-target hits.

**Figure 1.**
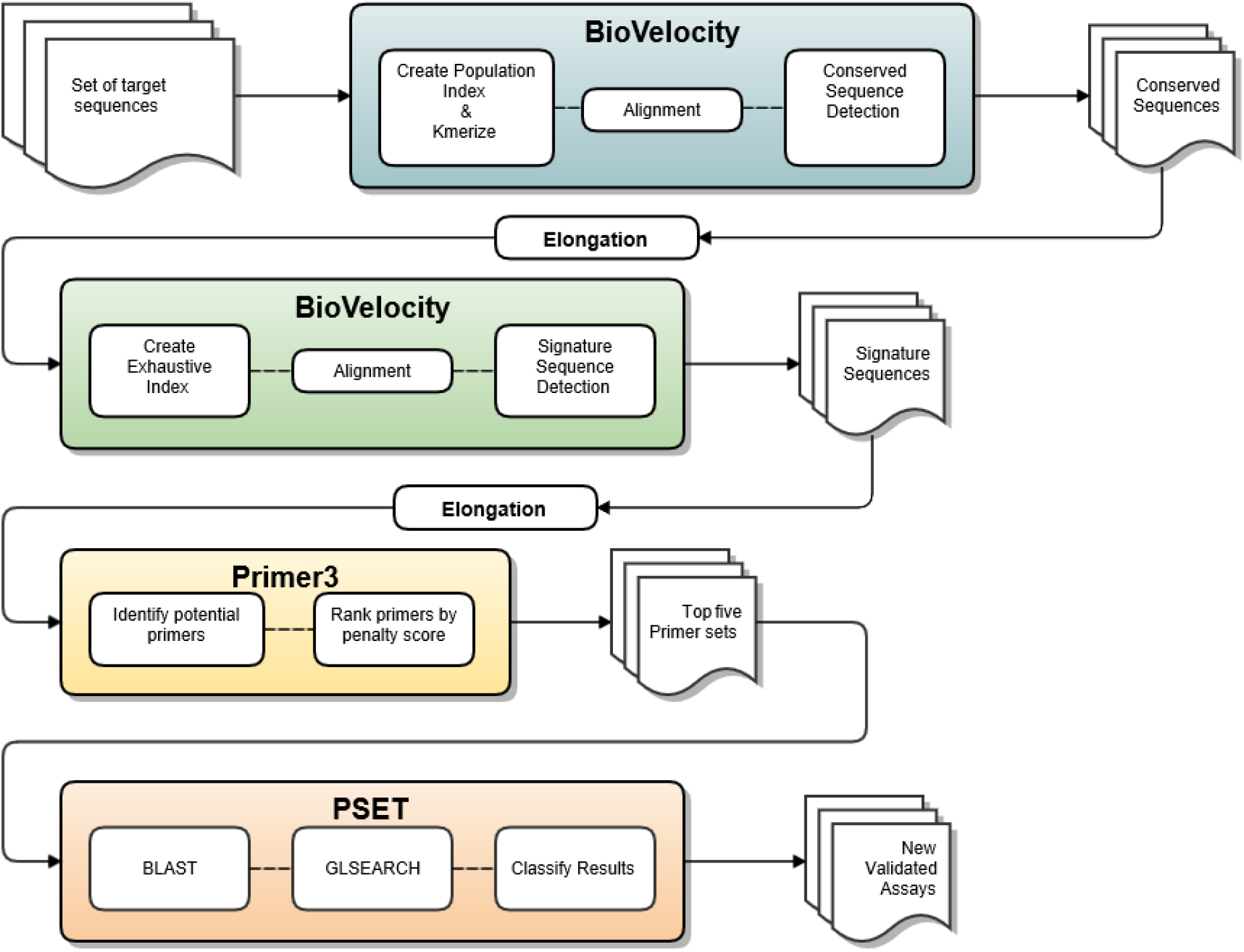
BioLaboro- New Assay Discovery pipeline.

### Performance assessment of currently available EBOV PCR assays on BOMV using PSET

The phylogenetic classification criteria for filoviruses were reported to be 64–77% similarity for species and 41–50% for genera, based on BLAST alignment results [37]. Compared to other ebolaviruses, the BOMV showed 55–59% nucleotide identity, 64–72% amino acid identity [20]. Given the sequence diversity, we assessed if the current diagnostic assays were effective against BOMV. We compared the current Ebola assay signatures against the three available complete genome sequences of BOMV (NCBI Accession numbers: MF319185.1, MF319186.1, and MK340750.1). Of the 30 Ebola assays examined (assay details in reference [32]), only two met our *in silico* criteria for successful detection (90% identity over 90% of the length for all primers and probe sequences in the amplicon). Of the two assays that did meet detection criteria for BOMV, neither had perfect matches to the primer sequences. The primer set match percentages (the average identity of all primers/probes) for all 30 Ebola assays against the three BOMV complete genomes are shown in the heat map (Table 1). The two assays which pass the *in silico* criteria are shown in green while the matches in red indicate failure or no alignment. Even the two assays that passed *in silico* criteria did not have perfect matches raising the possibility that these assays may fail in wet lab testing due to mismatches against currently available BOMV genomic sequences. Hence, as described below, we designed new assays using the BioLaboro platform.

**Table 1.** Heat map of PSET results of Ebola assays against BOMV sequences. Table 1 Legend: The average primer set percent identity matches for thirty current Ebola assays tested against the three BOMV genomes. Cells in green have an average primer identity over 90% which indicates likely primer binding while red cells have below 90% average identity or no alignment at all, indicating likely primer binding failure.

| Assay | MF319186.1 | MK340750.1 | MF319185.1 |
| --- | --- | --- | --- |
| EBO_GP | 94.21 | 94.21 | 94.21 |
| EBO1_2 | 72.50 | 75.00 | 72.50 |
| EBO3_4 | 74.34 | 74.34 | 74.34 |
| EBOGP | 78.41 | 76.75 | 78.41 |
| Ebola_Bundibugyo_MGB | None | None | None |
| Ebola_Bundibugyo_TM | None | None | None |
| Ebola_Ivory_Coast_MGB | None | None | None |
| Ebola_Ivory_Coast_TM | 72.22 | 69.44 | 70.83 |
| Ebola_Reston_MGB | None | None | None |
| Ebola_Reston_TM | None | None | None |
| Ebola_Sudan_MGB | 80.78 | 80.78 | 82.17 |
| Ebola_Sudan_TM | None | None | None |
| EboZNP | 75.66 | 76.01 | 75.66 |
| ENZ | 72.41 | 69.22 | 72.41 |
| Filo_AB | 73.48 | 73.48 | 73.48 |
| GAB_1 | 78.57 | 78.57 | 78.57 |
| KGH | None | None | None |
| Kulesh_MGB | 71.29 | None | 71.29 |
| Kulesh_TM | 74.74 | 74.74 | 74.74 |
| NGDS_Primary_amplicon | None | 61.39 | None |
| NGDS_Secondary_amplicon | 66.91 | None | 66.91 |
| pan_Ebola_Assay_MGB_EBOV | 76.74 | 76.74 | 76.74 |
| pan_Ebola_Assay_MGB_RESTV | 77.55 | 79.07 | 77.55 |
| pan_Ebola_Assay_MGB_SUDV | 70.63 | 72.30 | 70.63 |
| PanFiloL_1_2 | 85.55 | 85.55 | 85.55 |
| PanFiloL3_4 | 96.30 | 96.30 | 96.30 |
| Reston | 75.49 | 75.49 | 77.67 |
| Sudan | None | None | None |
| ZAI_NP | 80.59 | 80.59 | 80.59 |
| ZebovGP | None | None | None |

### Discovery of potential new BOMV assays using BioLaboro end-to-end pipeline

Using BioLaboro we ran a New Assay Discovery job to discover new BOMV signatures and determine their potential for accurate detection using PSET. In the first phase, BioVelocity was used to search for conserved and signature regions within the selected genomes. We selected the organism of interest by searching for “Bombali ebolavirus” from the database and selecting the three available complete genomes. The MF319185.1 genome was used as the algorithmic reference sequence as it is the same one that NCBI selected for the RefSeq database (Genbank ID: NC_039345.1). The algorithmic reference sequence was first split into k-mers of 50 bps each using a sliding window of 1 bp, which amounts to 18,994 k-mers to be evaluated with BioVelocity’s conserved sequence detection algorithm. BioVelocity found 27% (5,237) of these k-mers to be conserved in all three of the BOMV genomes. The conserved k-mers were then evaluated to determine overlapping segments and were combined into 120 conserved contigs. These contigs were next evaluated with BioVelocity’s signature sequence detection algorithm. The contigs were split into signatures with a max size of 250 bps (longer contigs were split into 250 k-mers with a step size of 1). The conserved k-mer sequences were evaluated against 563,843 complete genomes and plasmids from the NCBI GenBank repository. There were 291 k-mers sequences found to be signatures to BOMV. The signatures were then evaluated to determine overlapping reads and combined back into 119 signature contigs. Metrics for the BioVelocity run in phase one are shown in Table 2.

**Table 1.** BioVelocity run statistics for BOMV. Table 2 Legend. Total number of conserved and signature contigs along with total bases, average sequence length, median sequence length, and percent coverage of the BOMV genome

| Type | Contigs | Total Bases | Average Length | Median Length | Genome Coverage |
| --- | --- | --- | --- | --- | --- |
| Conserved | 120 | 11,117 | 93 | 81 | 58.3% |
| Signature | 119 | 11,046 | 93 | 81 | 58.0% |

In the second phase, Primer3 was used to identify potential primer pairs and probes for generating new PCR detection assays. For the BioLaboro pipeline, Primer3 is used to assess the suitability of a sequence to primer creation and generates viable primers which cover the identified signature regions. There were 151 primer sets created from the signatures run through Primer3 and assigned a penalty score to facilitate comparison of the results. The top five primer sets, by lowest penalty score (Table 3), were formatted and sent to the final step for validation using PSET.

**Table 3.**
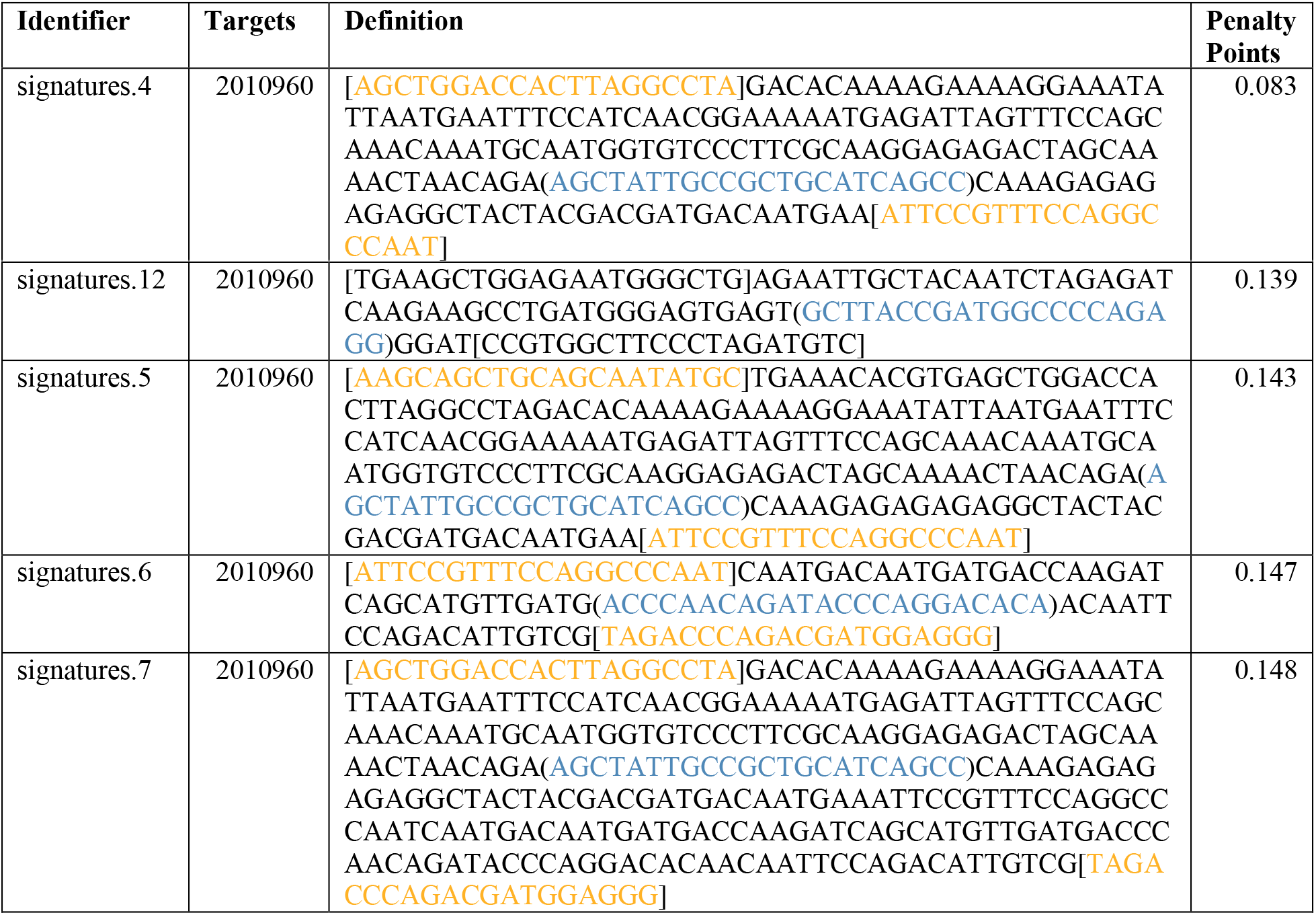
Primer3 identified PCR assays for BOMV. Table 3 Legend. The five new assays identified by Primer3 ranked by lowest penalty score. The Identifier column is an automated ID generated from the pipeline, the Targets column is the Taxonomy ID for BOMV, the Definition column contains the amplicon sequence with the primers in brackets (orange) and the probe in parentheses (blue), and the Penalty Points column contains the score generated after taking into account primer design parameters.

The signature regions identified by BioVelocity and the five new assays selected by Primer3 were mapped to the BOMV genome (Figure 2). As shown, the four assays are clustered in the nucleoprotein (NP) gene and one is in the glycoprotein (GP) gene. The signature segments (blue) indicate that there are many potential assay regions throughout the genome.

**Figure 2.**
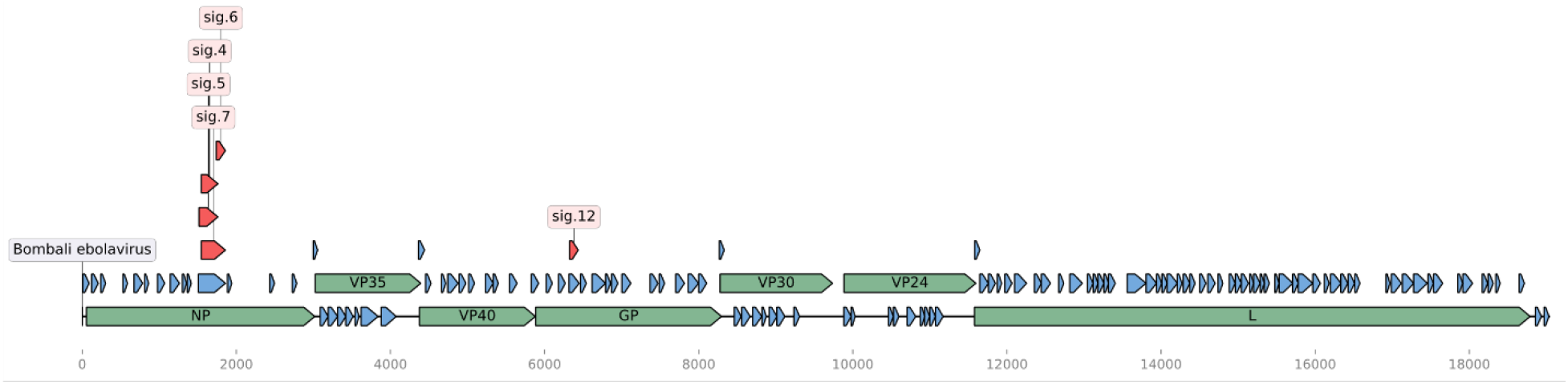
Linear map of BOMV genome. Figure 2 Legend: Linear map of the BOMV genome with annotations. The BOMV genes are colored in green, the signature segments are colored in blue, and the new assays are colored in red. The figure was created using the DNA Features Viewer Python library [38].

In the third phase, PSET was used to test the five newly designed assays identified by Primer3 *in silico* against publicly available sequences. PSET was used to analyze the primers and probes in the new assays using bioinformatics tools to identify potential false-positive and false-negative matches to NCBIs BLAST nucleotide sequence repositories (nt, gss, and env_nt) comprised of over 220 million sequences (as of last update in September 2019). BLAST+ was used to compare the assay amplicon sequences against these sequence repositories to identify matches. These matches were then used to create a custom library of sequences for GLSEARCH, a global-local sequence comparison tool in the FASTA suite of programs, which was used to search for the individual primers and probes. The resulting output was then processed and filtered based on pre-defined hit acceptance criteria. These criteria require that the assay components all hit to 90% identity over 90% of the component length, primer pairs were on opposite strands, and the total amplicon size was no greater than 1000 bps. The results were then validated by comparing the hits to the target NCBI Taxonomy identifier (ID), and true and false matches were reported. PSET results confirmed that the top five primer sets had true positive hits to all three BOMV genomes and no false positive hits to any other organism (Table 4).

**Table 4.** PSET results of BOMV assays. Table 4 Legend: True positive (TP: All assay components hit with >=90% identity over >=90% of the component length to the correct target), true negative (TN: Partial hit to assay amplicon but one or more assay components hit with <90% alignment to an incorrect target), false positive (FP: All assay components hit with >=90% identity over >=90% of the component length to an incorrect target), and false negative (FN: Partial hit to assay amplicon but one or more assay components hit with <90% alignment to the correct target) counts for each of the five new BOMV assays tested. The targets column is the NCBI Taxonomy ID of the target sequence, BOMV.

| Identifier | Targets | TP | TN | FP | FN |
| --- | --- | --- | --- | --- | --- |
| signatures.4 | 2010960 | 3 | 1647 | 0 | 0 |
| signatures.12 | 2010960 | 3 | 1678 | 0 | 0 |
| signatures.5 | 2010960 | 3 | 1678 | 0 | 0 |
| signatures.6 | 2010960 | 3 | 11 | 0 | 0 |
| signatures.7 | 2010960 | 3 | 1164 | 0 | 0 |

There are a high number of true negative (TN) results for 4 of 5 assays due to the similarity of the amplicon sequences with Zaire ebolavirus although these TNs are not expected to produce PCR positive results in wet lab experiments.

### Discovery of potential SARS-CoV-2 assays using BioLaboro end-to-end pipeline

Using BioLaboro, we ran a New Assay Discovery job to discover SARS-CoV-2 signatures and determine their potential for true positive viral detection. In the first phase, BioVelocity was used to search for conserved and signature regions within the selected genomes. Ninety six complete SARS-CoV-2 whole genome sequences were downloaded from the GISAID and uploaded to BioLaboro as a custom database. These 96 genomes were used as our reference set and EPI_ISL_404253 (Genbank ID: MN988713.1) was used as the algorithmic reference sequence. The algorithmic reference sequence was first split into k-mers of 50 bps each using a sliding window of 1 bp, which amounted to 29,833 k-mers to be evaluated with the conserved sequence detection algorithm. BioVelocity found 79% (23,542) of these k-mers to be conserved in all 96 of the SARS-CoV-2 genomes. The conserved k-mers were then evaluated to determine overlapping segments and were combined into 96 conserved contigs. These contigs were next evaluated with BioVelocity’s signature sequence detection algorithm. The contigs were split into signatures with a max size of 250 bps (longer contigs were split into 250 k-mers with a step size of 1). The conserved k-mer sequences were evaluated against 563,941 complete genomes and plasmids from the NCBI GenBank repository. There were 11,152 sequences found to be signature to SARS-CoV-2. The signatures were then evaluated to determine overlapping reads and combined back into 91 signature contigs. Metrics for the BioVelocity run in phase one are shown in Table 5.

**Table 5.** BioVelocity run statistics for SARS-CoV-2. Table 5 Legend: Total number of conserved and signature segments along with total bases, average sequence length, median sequence length, and percent coverage of the SARS-CoV-2 genome.

| Type | Contigs | Total Bases | Average Length | Median Length | Genome Coverage |
| --- | --- | --- | --- | --- | --- |
| Conserved | 96 | 28,246 | 294 | 221 | 94.5% |
| Signature | 91 | 27,888 | 306 | 241 | 93.0% |

In the second phase, Primer3 was used to identify potential primer pairs and probes for generating new PCR detection assays as described above for BOMV. There were 330 primer sets created from the signatures which were assigned a penalty score to facilitate comparison of the results. Primer sets were sorted by lowest penalty score and five potential assays were chosen manually in order to distribute potential candidates across the genome. These assays were then formatted and sent to the final step for validation using PSET. Primer sets sent to PSET are shown in Table 6.

**Table 6.**
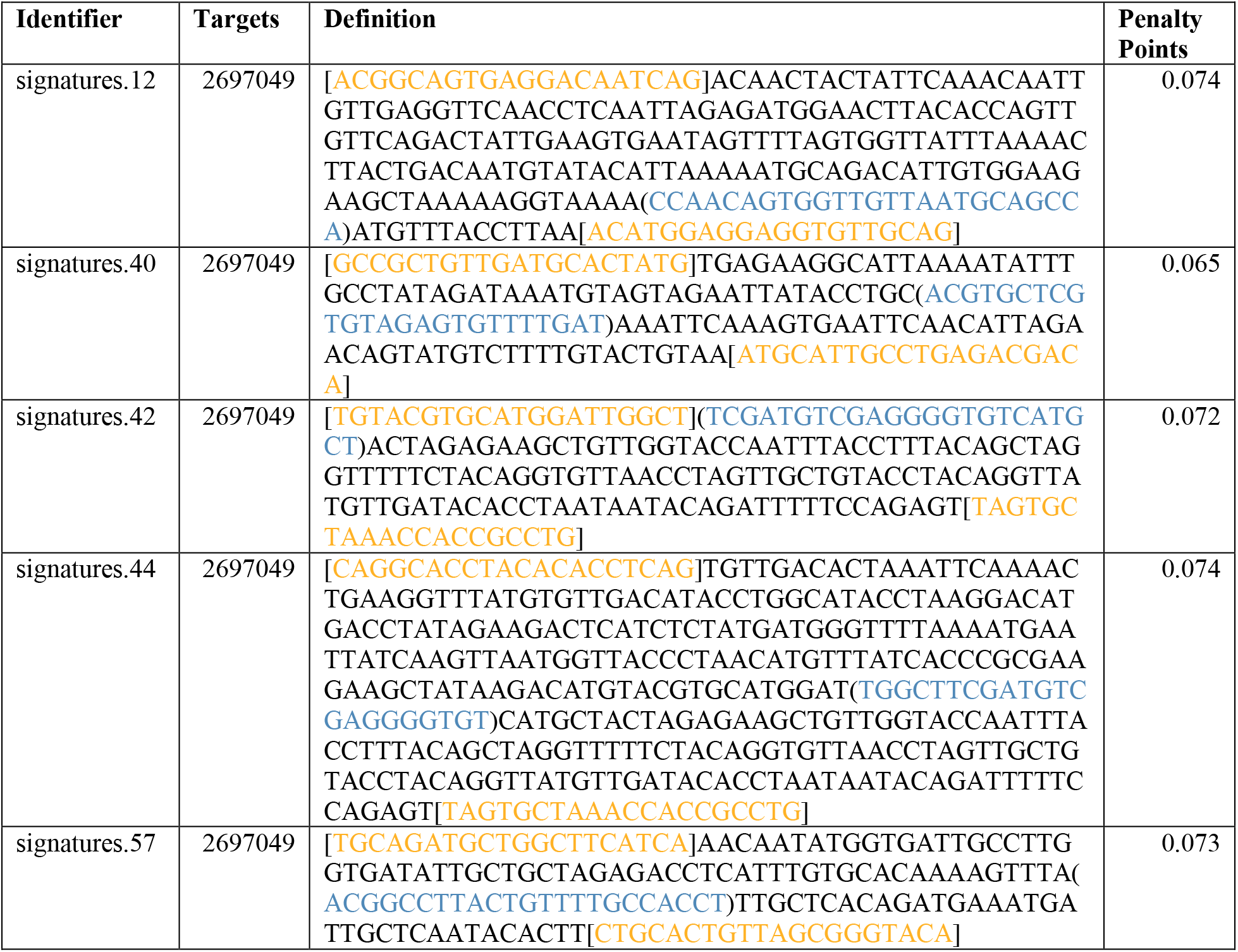
Primer3 identified assays for SARS-CoV-2. Table 6 Legend: The five new assays identified by Primer3 ranked by lowest penalty score. The Identifier column is an automated ID generated from the pipeline, the Targets column is the Taxonomy ID for SARS-CoV-2, the Definition column contains the amplicon sequence with the primers in brackets (orange) and the probe in parentheses (blue), and the Penalty Points column contains the score generated after taking into account primer design parameters.

The five new assays were mapped to the SARS-CoV-2 genome presented below (Figure 3). This figure shows that four assays are in the ORF1ab and one is located in the spike (S) gene. By comparison, the CDC and Corman group assays are clustered primarily at the 3’ end of the genome in the envelope (E) and nucleocapsid phosphoprotein (N) genes.

**Figure 3.**
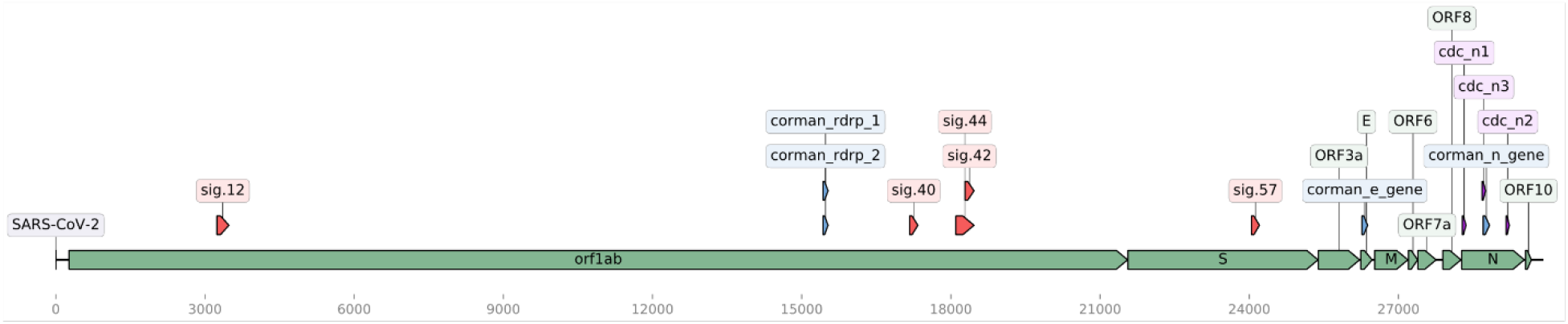
Linear map of SARS-CoV-2. Figure 3 Legend. A linear map of the SARS-CoV-2 genome with annotations. The genes and open reading frames (ORFs) are colored in green and the new assays are colored in red, Corman assays (blue), CDC assays (purple), and assays discovered in this study (red). The figure was created using the DNA Features Viewer Python library [36].

In the third phase, PSET was used to test the five newly designed assays identified by Primer3 *in silico* against publicly available sequences as described above for BOMV signatures. The results were then validated by comparing the hits to the target NCBI Taxonomy identifier (ID), and true and false matches were reported. PSET confirmed that the top five primer sets had true positive hits to all 145 (SARS-CoV-2) genomes (as of February 28, 2020) and no false positive hits to any other organism. An additional table lists the identifiers and metadata for these 145 complete genome sequences generated from human samples and not from other sources such as bat or pangolin [see Additional Table 2]. Results are shown alongside assays from CDC and Corman et al. [39, 40] (Table 7).

**Table 7.** PSET results of SARS-CoV-2 PCR assays. Table 7 Legend: True positive (TP: All assay components hit with >=90% identity over >=90% of the component length to the correct target), true negative (TN: Partial hit to assay amplicon but one or more assay components hit with <90% alignment to an incorrect target), false positive (FP: All assay components hit with >=90% identity over >=90% of the component length to an incorrect target), and false negative (FN: Partial hit to assay amplicon but one or more assay components hit with <90% alignment to the correct target) counts for each of the five new SARS-CoV-2 assays tested. The targets column is the NCBI Taxonomy ID of the target sequence, SARS-CoV-2.

| Identifier | Targets | TP | TN | FP | FN |
| --- | --- | --- | --- | --- | --- |
| signatures.12 | 2697049 | 145 | 2 | 0 | 0 |
| signatures.40 | 2697049 | 145 | 316 | 0 | 0 |
| signatures.42 | 2697049 | 145 | 275 | 0 | 0 |
| signatures.44 | 2697049 | 145 | 277 | 0 | 0 |
| signatures.57 | 2697049 | 145 | 359 | 0 | 0 |
| cdc_n1 | 2697049 | 145 | 284 | 0 | 0 |
| cdc_n2 | 2697049 | 145 | 282 | 0 | 0 |
| cdc_n3 | 2697049 | 145 | 14 | 273 | 0 |
| corman_e_gene | 2697049 | 144 | 4 | 283 | 1 |
| corman_n_gene | 2697049 | 145 | 22 | 266 | 0 |
| corman_rdrp_1 | 2697049 | 145 | 2957 | 420 | 0 |
| corman_rdrp_2 | 2697049 | 145 | 2853 | 52 | 0 |

There are a high number of true negative (TN) results for 4 of 5 assays due to the similarity of the amplicon sequences with SARS coronavirus near neighbors. The FP results from some Corman and CDC assays are due to near neighbor hits since these assays are pan assays. The one FN identified for corman_e_gene is to sequence EPI_ISL_410486 (Additional Table 2) which contains a large stretch of Ns over the majority of the amplicon sequence (80%) which is likely due to missing sequences.

We also tested the SARS-CoV-2 assays using PSET on near neighbor sequences that were generated during this outbreak, such as the bat and pangolin sequences. As expected the analyses showed a range of TP from pan assays, FN results due to sequence divergence (Table 8).

**Table 8.** PSET results of SARS-CoV-2 PCR assays against bat and pangolin SARS-CoV sequences. Table 8 Legend: TP, TN, FP and FN definitions are similar to Table 7.

| Identifier | Targets | TP | TN | FP | FN |
| --- | --- | --- | --- | --- | --- |
| signatures.12 | 2697049 | 0 | 0 | 0 | 8 |
| signatures.40 | 2697049 | 1 | 0 | 0 | 7 |
| signatures.42 | 2697049 | 3 | 0 | 0 | 5 |
| signatures.44 | 2697049 | 2 | 0 | 0 | 6 |
| signatures.57 | 2697049 | 1 | 0 | 0 | 7 |
| cdc_n1 | 2697049 | 3 | 0 | 0 | 5 |
| cdc_n2 | 2697049 | 1 | 0 | 0 | 7 |
| cdc_n3 | 2697049 | 1 | 0 | 0 | 7 |
| corman_e_gene | 2697049 | 8 | 0 | 0 | 0 |
| corman_n_gene | 2697049 | 2 | 0 | 0 | 6 |
| corman_rdrp_1 | 2697049 | 8 | 0 | 0 | 0 |
| corman_rdrp_2 | 2697049 | 3 | 0 | 0 | 6 |

### Discussion

Since its discovery in 2016, BOMV RNA has been detected in oral and rectal swabs as well as internal organs of *Mops condylurus* and *Chaerephon pumilus* bats in Sierra Leone, Kenya and Guinea [20, 41, 42]. These data add to the body of evidence suggesting that bats are a reservoir for filoviruses. However, these did not conclusively link the presence of viral RNA in bats to human infections with filoviruses. The discovery of BOMV in bats residing near and in human dwellings and residential areas further highlights the gaps in knowledge about ebolavirus diversity and ecology. Given these gaps and the human and economic impacts of ebolavirus disease, there is an ongoing need for ebolavirus biosurveillance and further characterization of BOMV. Availability of efficient viral RNA detection assays is critical for bio surveillance of these reservoirs.

Based on *in silico* analyses, we determined that current EBOV assays could potentially fail to detect BOMV sequences, and thus there is a need for BOMV specific assays. Using BioLaboro we rapidly designed and evaluated new, more specific assays. An advantage of BioVelocity is that the end user obtains results quickly with high confidence that the output of conserved and unique signature regions are accurate and not based on heuristics and probabilities. Additionally, Primer3 allows an end user to determine which signatures yield the best primers and probes, based on an objective penalty scoring system. PSET tests the PCR assays *in silico* against the latest versions of public sequence repositories, including newly added strain genomes, to validate that the primers and probes match only to their intended targets. BioLaboro’s easy-to-use Graphical User Interface (GUI) provides full functionality for submitting a new job, viewing the current job queue, checking results from previously completed jobs, and exploring the system database management and settings. The dedicated large RAM system easily supports multiple users with discrete logins and rapid operations. Moreover, the user-friendly GUI allows scientists without command-line experience to design and evaluate an assay for immediate wet lab testing.

Due to the relative novelty of BOMV there are currently only three complete genomes available from NCBI. As more samples are identified and sequenced the genetic diversity will likely increase. During the 2014 - 2016 Western African EBOV outbreak, rapid accumulation of inter- and intra-host genetic variations were observed [43]. Since many of the nucleotide mutations altered protein sequences, it became apparent that the changes should be monitored for impacts on diagnostics, vaccines, and therapies critical to outbreak response. In an earlier study conducted to decipher the impact of the then-available diagnostics, we determined that many of the real-time reverse transcription PCR (rRT-PCR) assays that were in use during that outbreak identified regions outside of those that BioVelocity selected as unique to EBOV, SUDV, and RESTV [32]. In another study, signature erosion of diagnostic assays was identified during the 2018 outbreak in North Kivu and Ituri Provinces of the Democratic Republic of the Congo [44]. Using the *in silico* methods described here, only two of the 30 EBOV rRT-PCR assays evaluated against BOMV target sequences met our criteria for successful detection and neither showed perfect matches. For ongoing biosurveillance, we recommend wet lab testing and validation of the rRT-PCR assays described here to ensure detection of BOMV.

We also tested the BioLaboro pipeline with available SARS-CoV-2 viral genomes and rRT-PCR assays. We identified five signature sequences distributed across the genome and in different regions than those of the CDC or German group [39, 40]. There are seven assays currently in use (3 CDC and 4 German) for SARS-CoV-2 diagnostics and they all produced true positive results against the human-derived sample sequences without any signs of signature erosion. A few whole genome sequences that showed false negative results were from environmental samples (Bat and Pangolin origin) indicating that the diagnostic assays are specific for human isolates. The assays that produced true positive results were pan assays. The lack of signature erosion is in agreement with the whole genome sequence data analyzed thus far (February 28, 2020). A simple pipeline consisting of Multiple Sequence Alignment using Fast Fourier Transform (MAFFT), snp-sites, and R we calculated 228 single nucleotide polymorphisms (SNPs) (159 unique) across 145 genomes [45-47]. Each of the 145 WGS contains less than 10 SNPs with the exception of one having 25. None of the variations impact the diagnostic assay signatures. However, real-time monitoring of these assays against WGS as they become available, will enable rapid identification of signature erosion if it occurs and generation of new assays as needed. The newly designed assays we have described here need to be validated in wet lab testing and with appropriate clinical matrices to determine their performance. However, we have demonstrated that the BioLaboro pipeline can be used effectively and rapidly to validate available assays and to design new assays using genome sequences of newly emerging pathogens.

### Conclusions

By periodically re-running BioLaboro on emerging and reemerging pathogen sequences as they become available, over time the relative diversity can be monitored, and assays can be updated to remain current with regards to available data. By tracking assay performance measures over time, one can evaluate the efficacy of MCMs on a routine basis. These analyses would ensure that the most accurate MCMs are available when an outbreak response is necessary. In this study we demonstrate the value of real-time genomic sequencing and MCM evaluation to provide actionable information before and during a public health emergency. Combined with an active biosurveillance of zoonotic reservoirs and generation of sequence data to understand the genetic diversity of these pathogens, BioLaboro is broadly applicable for providing effective diagnostics and medical countermeasures during a crisis involving future threats.

## Methods

### BioLaboro System Description

BioLaboro is an application for rapidly designing *de novo* assays and validating existing PCR detection assays. It is composed of three tools: BioVelocity, Primer3, and PSET which are built into a pipeline for user-friendly new assay discovery via an interactive graphic user interface.

**BioVelocity** is a bioinformatics tool based on an innovative algorithm and approach to genomic reference indices [32]. Using a fast and accurate hashing algorithm, BioVelocity can quickly align reads to a large set of references. BioVelocity takes advantage of large RAM systems (hardware specification described in Additional Table 1) and creates a k-mer index of all selected reference sequences (e.g. GenBank) by identifying all possible base pair sequences of various k-mer lengths. This index is used to determine all possible matches between query sequences and references, simultaneously. The advantage of this approach is that it allows for rapid identification of sequences conserved within or omitted from a set of target references. Thus, the used has high confidence in the conserved and signature (unique) designations because they are not based on heuristics and probabilities.

**Primer3** [48] is a tool for designing primers and probes for real-time PCR reactions. It considers a range of criteria such as oligonucleotide melting temperature, size, GC content, and primer-dimer possibilities. Potential new primer sets are identified within the signature regions using Primer3 analysis. For the BioLaboro pipeline, Primer3 has been configured to analyze 22 parameters influencing suitability of a sequence to primer creation and construct viable primers which cover identified signature regions. These primers are scored with a penalty scoring system to attempt to determine the fitness of the resulting primers thus allowing an end user to assess which signatures yield the best primers when the signatures themselves may be of similar size.

**PSET** is configured to test PCR assays *in silico* against the latest versions of public sequence repositories to determine if the primers and probes still match only to their intended targets. An elaborate description of PSET is provided in ref [32]. As NCBI’s database and other public databases are updated periodically, newly added genomic sequences can reveal where primers and probes may no longer be functional or where PCR assays may detect previously un-sequenced near neighbors. Using this information, an assay provider can be better aware of potential false hits and design new primers when false hits become an issue. PSET is used to test currently deployed assays as well as new assays designed using BioLaboro’s capabilities.

The BioLaboro application is composed of a fully functional GUI front-end that allows users to submit jobs to the back-end bioinformatics pipeline hosted on a dedicated large RAM system. The system has multi-user capability with discrete logins and a single job queue. The landing screen, shown in Figure 4, gives the user options for submitting a new job, viewing the current job queue, checking results from previously completed jobs, and exploring the system database management and settings.

**Figure 4.**
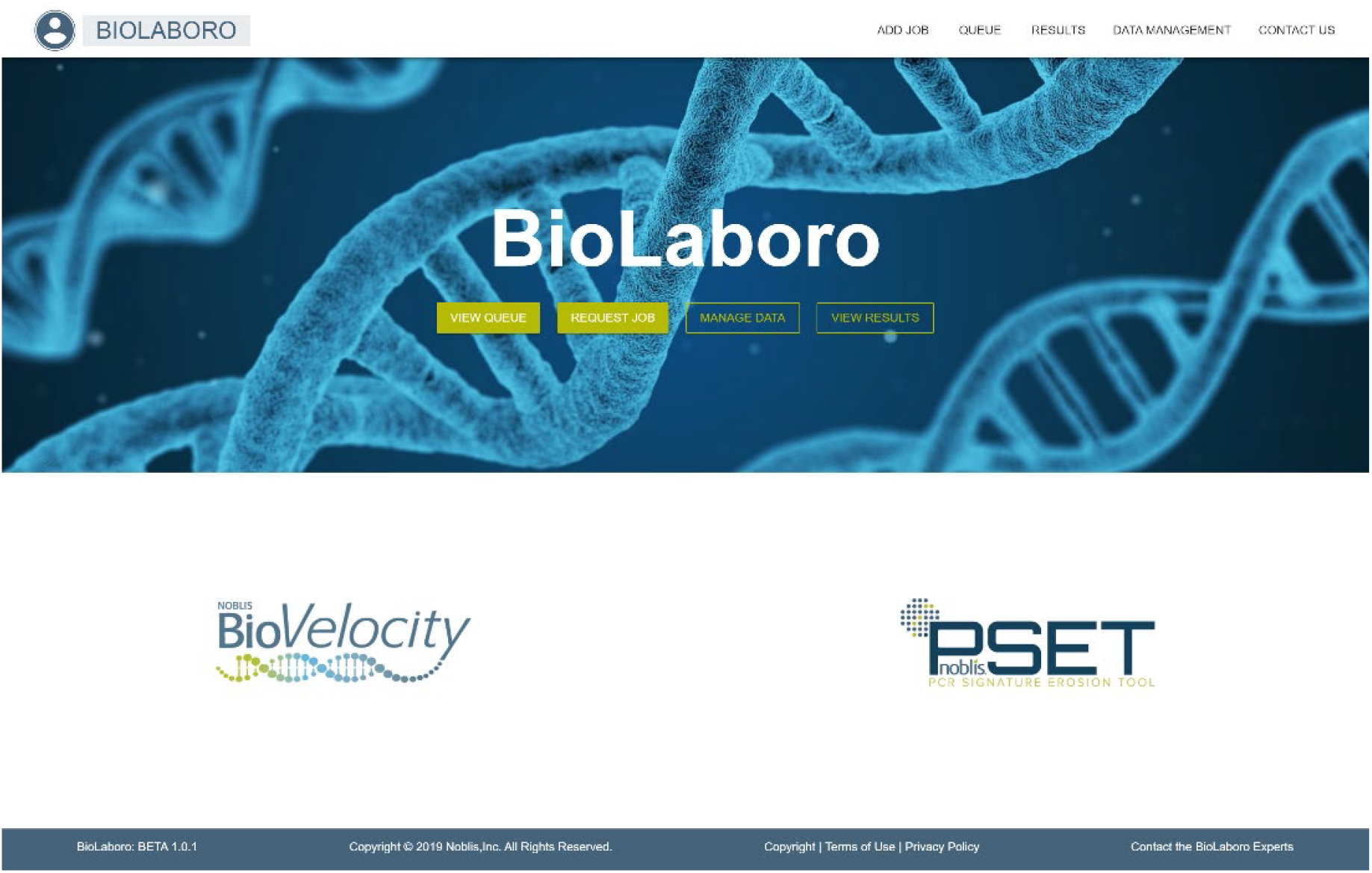
The landing page of BioLaboro.

The BioLaboro application allows job submissions through a simplified user interface designed for scientists with minimal or no command-line experience. The user can search for sequences of interest using the built-in Organism Select tool, Figure 5, which allows for searching on free text, NCBI Accession number, or NCBI Taxonomy ID. The results can then be filtered using a “smart filter” which will only include sequences within +/- 10% of the calculated median genome length of the results. This tool is useful for automatically excluding plasmids or sequence fragments which can negatively impact signature identification. Alternatively, custom sequence size filters can also be used if the user wants to target specific plasmids or chromosomes. Once all sequences are selected and added the user can optionally choose a specific sequence to serve as the algorithmic reference.

**Figure 5.**
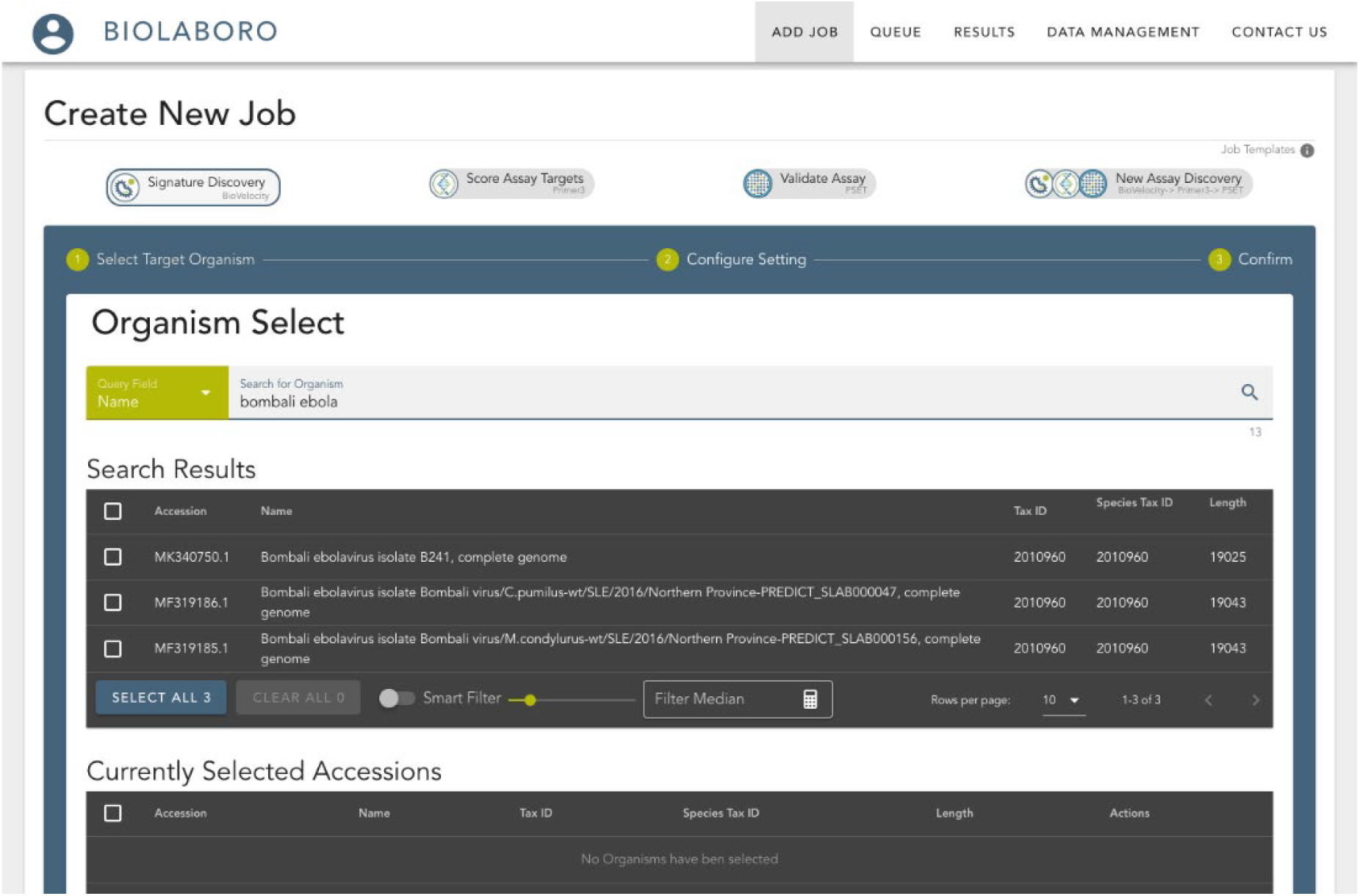
The Job Submission page for the BioVelocity component of BioLaboro showing the BOMV sequences. BioLaboro employs a queuing system to manage job submissions due to the high computational requirements of the BioVelocity algorithm. The queue page identifies the currently running job, the ordered list of queued jobs, and a list of previously finished jobs with timestamps and completion status, Figure 6. Each finished job can be re-launched from this dialog in the future with previously used parameters while utilizing the newest available datasets.

**Figure 6.**
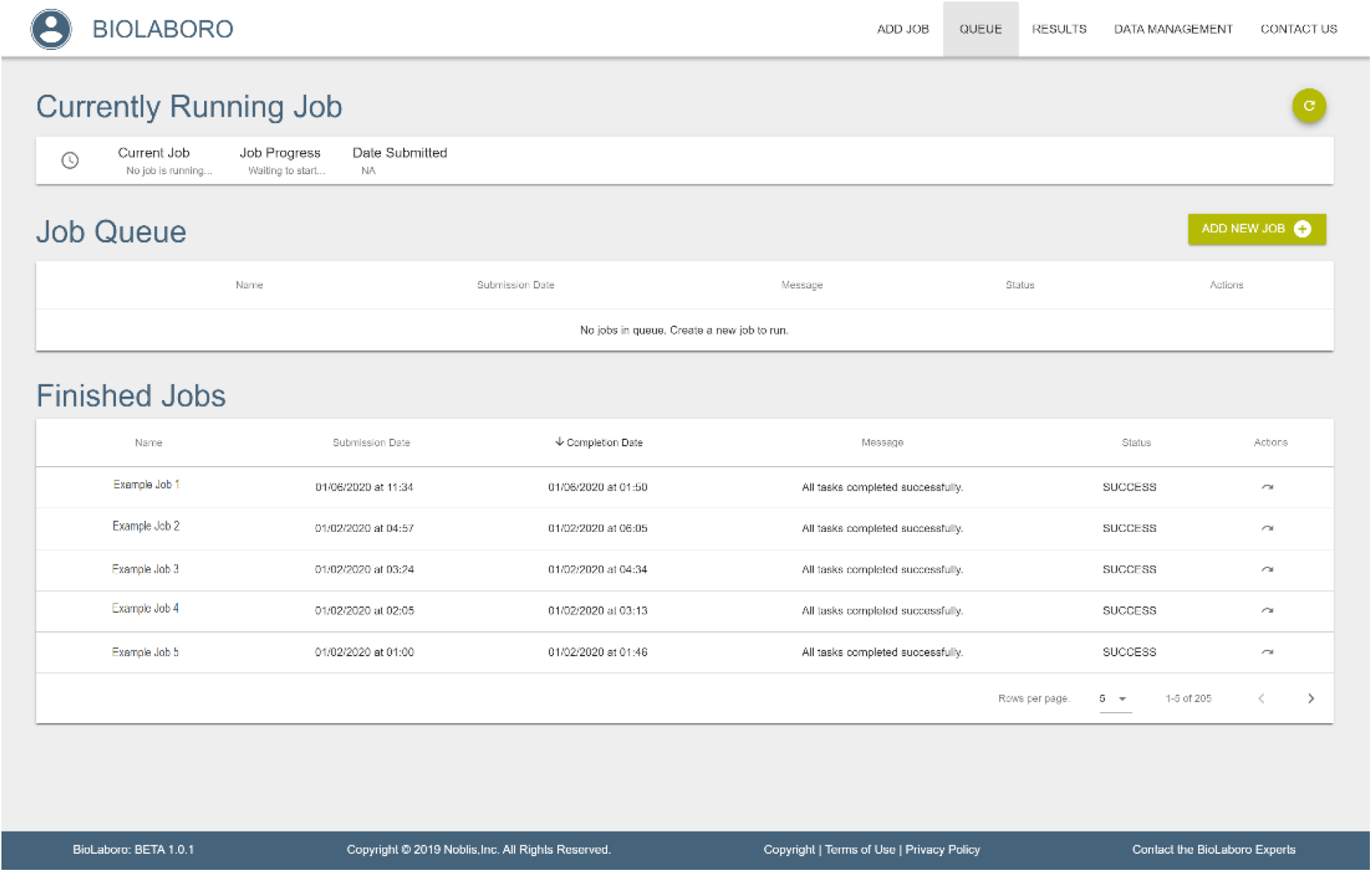
The BioLaboro job queue. The queue includes three sections: 1) The currently running job is identified with information on the current sub-process, 2) The job queue is shown, which can be re-arranged as needed, and 3) The finished jobs section shows completed jobs with timestamp metadata and an option to re-run the same job with identical parameters.

The Database Management and Settings Page contains many options for maintaining system data and pipeline settings. BioLaboro’s simple user interface design allows the user to manage the genomic reference databases from within the GUI. The GenBank Repository page, Figure 7, lists the databases loaded on the system along with the number of sequences each contains. These databases are used for selecting sequences for signature creation and can be updated on-demand by submitting a sync job to the queue. Other GenBank databases or custom sequence databases can also be added and maintained through this entry point.

**Figure 7.**
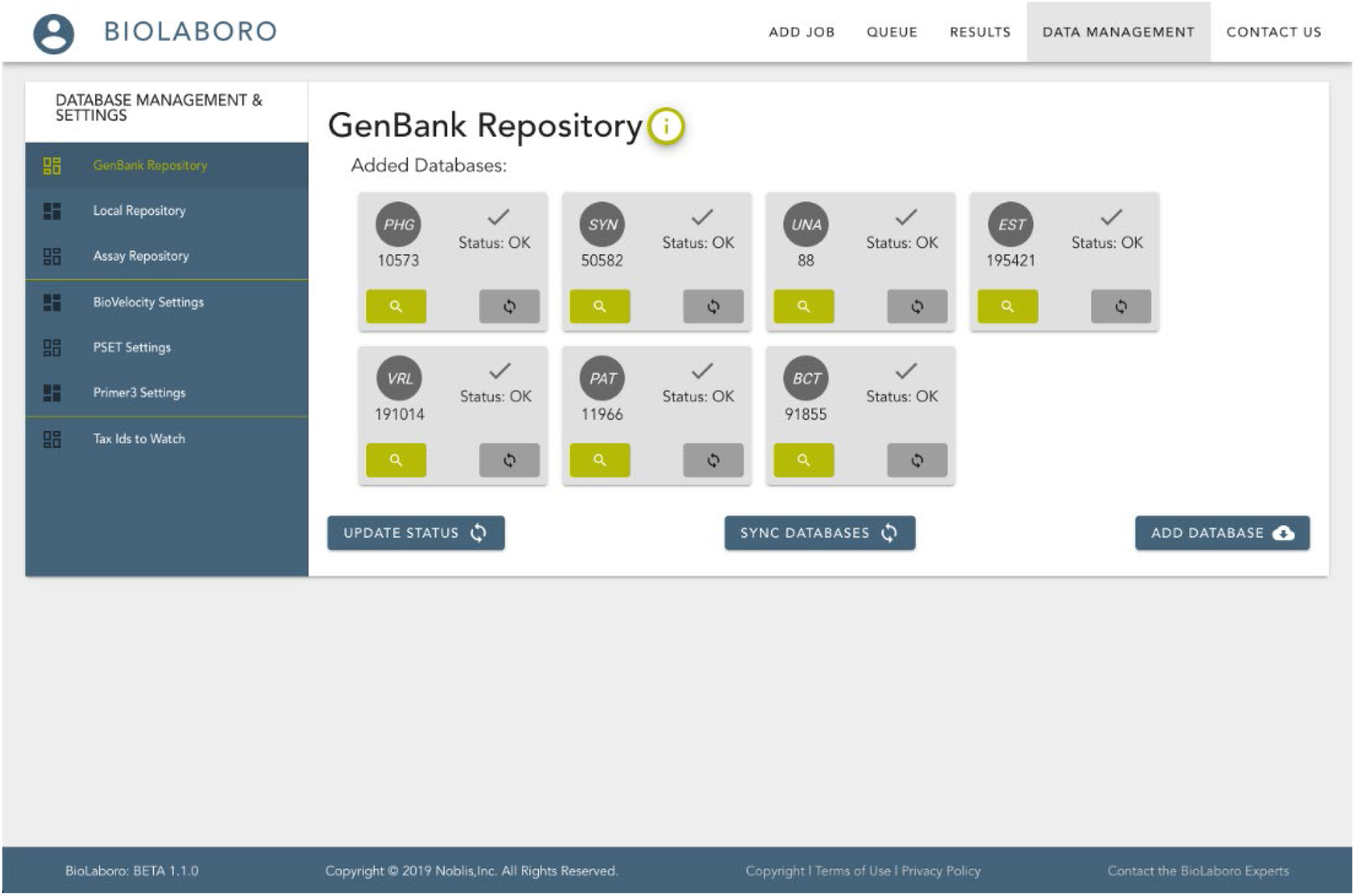
The BioLaboro GenBank Repository located in the Database Management and Settings page.

PCR detection assays are also maintained in the system and can be viewed and searched for in the Assay Repository. The system maintains a history of past assay runs so that the user can track performance of assays over time, as well as re-run assays when new sequences of interest are available. Lastly the Database and Management Settings page maintains the default settings for each of the three system tools which can be permanently set here, or specifically tailored for each job at run-time. Results for completed runs are available from the Results page which allows the user to view raw data or system generated reports as well as download them to local copies.

## Supporting information

Mitchell_et_al_BioLaboro_Sup_table

## Supplementary Information

**Additional Table 1.** Large RAM System Hardware Specifications.

| Component | Value |
| --- | --- |
| Architecture | ppc64le |
| RAM | 1 terabyte |
| CPUs | 160 |
| Threads per core | 8 |
| Cores per socket | 5 |
| Sockets | 4 |
| Local Storage | 15 terabytes |

Additional Table 2. (gisaid_sequences.xlsx) List of whole genome sequences downloaded from GISAID and used in this study. The “Signature Creation” column indicates which sequences were used to generate the signatures, and the “Validation” column indicates which sequences were used to validate the signatures with PSET.

## Declarations

**Ethics approval and consent to participate:** Not applicable

**Consent for publication**: Not applicable

**Availability of data and materials:**

Weekly updates of SARS-CoV assay performance using PSET are posted at Viological.org. (http://virological.org/t/preliminary-in-silico-assessment-of-the-specificity-of-published-molecular-assays-and-design-of-new-assays-using-the-available-whole-genome-sequences-of-2019-ncov/343/16).

The datasets analyzed during the current study are available in the following:

Three available complete genome sequences of BOMV NCBI Accession numbers: MF319185.1 https://www.ncbi.nlm.nih.gov/nuccore/MF319185.1/, MF319186.1 https://www.ncbi.nlm.nih.gov/nuccore/MF319186.1, MK340750.1 https://www.ncbi.nlm.nih.gov/nuccore/MK340750.1

Complete genome sequences of SARS-CoV-2: We gratefully acknowledge the authors, originating and submitting laboratories of the sequences from GISAID’s EpiFlu™ Database on which this research is based. https://www.gisaid.org/

NCBI Taxonomy database: https://www.ncbi.nlm.nih.gov/taxonomy

EMBL-EBI FASTA GLSEARCH: https://www.ebi.ac.uk/Tools/sss/fasta/nucleotide.html

## Authors’ contributions

MH, DN, SM, SS conceptualized the study. MH, DN, SM, ND, MI, ST developed BioLaboro. MH, DN, SM, SS analyzed the data. BG provided study resources. MH, DN, SM, MI, TB, KJ, SS wrote and edited the manuscript.

## Acknowledgements

The views expressed in this article are those of the authors and do not necessarily reflect the official policy or position of the DBPAO, JPEO-CBRND, Department of Defense, the US Government, nor the institutions or companies affiliated with the authors. BG is a US Government employee and this work was prepared as part of his official duties. Title 17 of the United States Code §105 provides that “Copyright protection under this title is not available for any work of the United States Government.” Title 17 of the United States Code §101 defines a US Government work as a work prepared by a military service member or employee of the US Government as part of that person’s official duties. We gratefully acknowledge the Authors and the Originating and Submitting Laboratories for the SARS-CoV-2 sequences and metadata shared through GISAID (https://www.gisaid.org/) on which SARS-CoV-2 part of this study is based.

## Abbreviations

bp: base pair (bps, plural base pairs)
BDBV: Bundibugyo ebolavirus
BLAST: Basic Local Alignment Search Tool
BOMV: Bombali ebolavirus
CFR: Case Fatality Ratio
COVID-19: Coronavirus Disease 2019
DNA: deoxyribonucleic acid
DRC: Democratic Republic of the Congo
EBOV: Zaire ebolavirus
EMA: European Medicines Agency
EVD: Ebola Virus Disease
FDA: Food and Drug Administration
FN: False Negative
FP: False Positive
GC: guanine cytosine
GP: glycoprotein
GUI: Graphic User interface
ID: identifier
mAb: monoclonal antibody
MAFFT: Multiple Sequence Alignment using Fast Fourier Transform
MCM: medical countermeasure
N: nucleocapsid phosphoprotein
N/A: Not available
NCBI: National Center for Biotechnology Information
NIAID: National Institute of Allergy and Infectious Diseases
NP: nucleoprotein
NPC1: Niemann-Pick C1
ORF: Open Reading Frame
PALM: Pamoja Tulinde Maisha study
PCR: polymerase chain reaction
PSET: PCR signature erosion tool
RAM: random access memory
RESTV: Reston ebolavirus
RNA: ribonucleic acid
rRT-PCR: real-time reverse-transcription polymerase chain reaction
rVSV: recombinant vesicular stomatitis virus
SARS: severe acute respiratory syndrome
SNP: single nucleotide protein
SUDV: Sudan ebolavirus
TAFV: Taï Forest ebolavirus
TN: True Negative
TP: True Positive
WGS: Whole Genome Sequence
WHO: World Health Organization

